# CD99 regulates cancer cell transendothelial migration and endothelial cell function via CDC42 and actin remodelling

**DOI:** 10.1101/760934

**Authors:** Aarren J. Mannion, Adam F. Odell, Alison Taylor, Pamela F. Jones, Graham P Cook

**Affiliations:** Leeds Institute for Medical Research, University of Leeds School of Medicine, St. James’s University Hospital, Leeds LS8 2BH, UK

**Author notes:** Department of Oncology and Pathology, Karolinska Institute, Stockholm, Sweden (Aarren J. Mannion). School of Health Sciences, York St. John University, York YO31 7EX, UK. These authors contributed equally to this work (Pamela F. Jones, Graham P Cook). **Correspondence** to Graham Cook.

**Keywords:** CD99, CDC42, transendothelial migration, metastasis, breast cancer, extravasation, cytoskeleton

## Abstract

Metastasis requires tumour cells to cross endothelial cell (EC) barriers and this occurs using mechanisms similar to those used by extravasating leucocytes during inflammation. The cell surface receptor CD99 is expressed by leucocytes and EC and participates in inflammatory transendothelial migration (TEM). CD99 is also expressed by tumour cells and we have analysed its role in tumour progression and cancer cell TEM. In a xenograft model, CD99 expression inhibited the metastatic progression of human breast cancer. *In vitro*, tumour cell CD99 was required for adhesion to ECs. However, tumour cell CD99 inhibited the invasion of the endothelial barrier by breast and prostate cancer cells and TEM itself. Furthermore, tumour cell CD99 depletion was associated with cytoskeletal remodelling. Loss of EC CD99 enhanced endothelial barrier function and reduced tumour cell TEM. Mechanistically, CD99 loss enhanced the expression and activity of CDC42, a known cytoskeletal organiser. CDC42 positively regulates EC angiogenic activity and the enhanced CDC42 activity resulting from loss of EC CD99 increased angiogenesis. As a signal transduction hub, CDC42 activity impacts upon many of the hallmarks of cancer. The functional link between CD99 and CDC42 identified here implicates CD99 in regulating these diverse pathways by modulation of CDC42 activity.

## Background

Ninety percent of cancer deaths are attributable to metastatic progression [1]. Key steps in the metastatic cascade include the interactions occurring between the disseminating, metastatic cancer cells and the endothelium [1]. Migrating cancer cells intravasate in order to gain access to the lymphatic and haematogenous vasculature [2] and circulating cancer cells which survive anoikis, exposure to shear flow and immune surveillance, travel to distant sites where they extravasate into secondary tissues [1]. Adhesion, intravasation and extravasation all require intimate interactions between cancer cells and endothelial cells (EC), with both cell types contributing to transendothelial migration (TEM) [3]. Leukocytes undergo TEM when extravasating from blood to infected tissues as part of the inflammatory process and this mechanism provides a model for how cancer cell TEM might operate during metastasis [4, 5].

The type 1 transmembrane receptor CD99 is expressed in a variety of tissues including haematopoietic lineage cells and EC [6]. Amongst its reported functions, CD99 regulates the adhesion and TEM of haematopoietic cells [7, 8]. In the endothelium, CD99 resides at the EC borders in a complex with ezrin, soluble adenylyl cyclase (sAC) and protein kinase A (PKA) (ref. 9). Endothelial cell CD99 is also localised within the lateral border recycling complex (LBRC), an intracellular compartment contiguous with the plasma membrane and located proximal to endothelial cell-cell junctions [9, 10]. During TEM, CD99 undergoes homophilic interactions between the leukocyte and endothelium which facilitates the movement of the leukocyte between the EC junctions [7, 9]. This engagement of endothelial CD99 triggers sAC, which activates PKA and enables the rapid turnover of CD99 from the LBRC to the endothelial cell junction to facilitate transmigration [9]. Although well established in leucocyte TEM, the involvement of CD99 in cancer progression remains enigmatic [11]. It is highly expressed in several haematopoietic malignancies as well as solid tumours, in particular in Ewing’s sarcoma where lower CD99 expression results in reduced migration, tumour growth and metastasis [12, 13]. However, in other cancers (such as osteosarcoma and gastric cancer), decreased CD99 expression is associated with tumour progression [11].

The TEM of leucocytes requires cytoskeletal rearrangements in both the migrating cell and the endothelium. Members of the Rho GTPase family control many cellular responses, including cytoskeletal organisation and cellular migration [14, 15]. Hence, Rho GTPase activity is implicated in many aspects of tumour and vascular biology, including maintenance of the endothelial barrier [16, 17], development of the vasculature during angiogenesis [18], and in facilitating cancer cell dissemination [19]. The CDC42 molecule is one of several Rho GTPases known to play a role in both vascular development and cancer metastasis [20]. Like other Rho GTPases, CDC42 cycles between an active GTP bound form and an inactive GDP bound form. This cycling is regulated by guanine exchange factors (GEFs) and GTPase activating proteins (GAPs) which activate and inactivate Rho GTPases respectively [15]. In its active, GTP-bound confirmation, CDC42 acts together with N-WASP to activate the ARP2/3 complex, facilitating the nucleation of actin and regulating cytoskeletal organisation [21, 22].

We have dissected the role of CD99 in the adhesion and TEM of metastatic cancer cells. We show that CD99 regulates metastatic progression *in vivo* and that cytoskeletal rearrangements determine cancer cell TEM *in vitro*. Furthermore, EC CD99 controls endothelial barrier function, thereby regulating tumour cell TEM. We show that CD99 mediates these effects by negatively regulating CDC42 activity. These data identify CD99 as a regulator of cytoskeletal organisation, with impact on tumour migration and TEM, key components of the metastatic pathway.

## Methods

### Cells and cell culture

Human umbilical vein endothelial cells (HUVEC) and Human Dermal Fibroblasts (HDF) were purchased from Promocell. MDA-MB-231 and MCF7 cells were purchased from the European Collection of Cell Cultures (ECACC). Brain metastatic derived MDA-MB-231 cells (BrM) were a kind gift from Dr. Mihaela Lorger (University of Leeds) and the SKBR3 breast cancer cell line was a kind gift from Dr. Laura Matthews (University of Leeds). PC3 cells were purchased from the ECACC. EC lines were cultured using Endothelial Cell Basal Medium (ECBM; from Promocell), supplemented with 2% FCS (v/v), 0.4% endothelial cell growth supplement, 0.1ng/ml epidermal growth factor (recombinant human), 1ng/ml basic fibroblast growth factor (recombinant human), 90μg/ml heparin, 1μg/ml hydrocortisone. Cells were grown on 0.2% gelatin (Sigma) (w/v in PBS) coated plates. HUVEC cells were grown to passage 5 or 6. MDA-MB-231 and MDA-BrM breast cancer cell lines were cultured in 10% (v/v) foetal calf serum (FCS; Sigma) supplemented RPMI-1640 (Sigma) and required passaging every 3-5 days. HDF, MCF7, SKBR3 and PC3 cells were cultured in 10% FCS supplemented Dulbecco’s modified essential media (DMEM; Sigma). All cell lines were incubated at 37°C under 5.0% CO_2_.

### Flow cytometry

Cultured cells were PBS washed and trypsinised with 1x Acutase (Gibco). Cells were washed in ice cold PBS followed by centrifugation at 300 x *g* for 5 min. After repeated washing in PBS, cells were resuspended in 100μl FACS buffer (PBS, 2% FCS, 0.09% NaN_3_) and stained with fluorophore conjugated antibodies (CD99-APC, HCD99 12E7 from Biolegend) and relevant isotype control antibodies at 1×10^6^ cells per 100μl staining buffer for 30 min at RT. Stained cells were washed and fixed in Cytofix Fixation buffer (BD Biosciences) before analysis using a LSRII flow cytometer (BD Biosciences).

### Tumour-EC adhesion assay

EC were seeded at a density of 1×10^4^ per well of a 96 well plate (Corning) and incubated for 24-48 h or until confluent monolayers were present in each well. MDA-MB-231 cells were labelled with Cell Tracker Green for 30 min in serum free RPMI (SFM-RPMI) media at 37°C. MDA-MB-231 cells were washed in SFM RPMI once before being seeded at 1×10^4^ per confluent EC monolayer. Adhesion assay was incubated at 37°C and MDA-231 cells were allowed to adhere to the EC monolayers for 15, 30, 60 and 120 minutes, after which each plate was washed 1x in PBS, and fixed in 4% (w/v) paraformaldehyde (PFA)(Sigma) for 10 minutes, and washed twice in PBS before storing at 4°C and analysed using an Incucyte Zoom Live Cell Imager (Essen bioscience).

### RNA interference

HUVEC and MDA-MB-231 were transfected with SMARTpool siRNA (Dharmacon) targeting CD99. Control siRNA was used alongside untransfected (or mock transfected) cells. Transfections were performed using Lipofectamine 200 RNAiMax (Invitrogen) transfection agent and Opti-MEM I Reduced Serum Medium, GlutaMAX Supplement (Gibco) according to manufacturer’s instructions. Cells were transfected with 30pmol siRNA in a 6 well plate (2-4×10^5^ cells/well) and scaled accordingly. Briefly, for a single well of a 6 well plate, 30pmol siRNA duplexes were made in 250μl of OptiMEM media and incubated at RT for 5 min. At the same time 5μl Lipofectamine was made up in 250μl of OptiMEM and incubated at RT for 5 min. siRNA and Lipofectamine mix were combined within the 6 well plate and gently mixed before incubating at RT for 20 minutes. Following this OptiMEM suspended cells were added to siRNA Lipofectamine complexes at 2-4×10^5^ cells in 1ml of OptiMEM. Cells were incubated in this mixture for 4-6 hours, before transfection medium was aspirated and replaced with supplemented normal culture medium. siRNA treated cells were incubated for 24-72 hours before use in downstream assays. ON-TARGETplus Human CD99 (4267) siRNA – SMARTpool (5’-GGAUGGUGGUUUCGAUUUA-3’, 5’-CUUCAUUGCUUACCAGAAA-3’, 5’-GAACCCACCCAAACCGAUG-3’, 5’-CGUUUCAGGUGGAGAAGGA-3’); Control (ON-TARGETplus Non-targeting control pool, Dharmacon).

### Western blotting

Cells were washed in PBS before being lysed in lysis buffer (2% SDS (Sigma) in PBS). Cell lysates were boiled at 100°C for 5 minutes, and sonicated using Sonicator (Soniprep 150 – MSE) for 1-2 seconds. Protein concentration of samples was then determined using Pierce™ bicinchoninic acid assay BCA Protein Assay Kit (Thermo Scientific). Cell lysates were then mixed with 2x sample buffer (100 mM Tris-HCl, 4% SDS, 20% glycerol, 0.2% bromophenol blue, 10% β-mercaptoethanol) and stored at −20°C before being loaded into 8, 10, 12 or 15% SDS page gels (8,10, 12 or 15% acrylamide, 380mM Tris pH8.8, 0.1% SDS, 0.1% APS, 0.1% TEMED) set with 4% stacking gel (4% acrylamide, 120mM Tris pH6.8, 0.1% SDS, 0.1% APS, 0.1% TEMED) for electrophoresis. Samples were loaded along with SeeBlue Plus 2 pre-stained protein standard (Invitrogen) and were electrophoresed for 1h 20min at 120V (12V/cm). Following transfer, membranes were blocked and probed with primary antibodies (CD99, 12E7, Biolegend; CDC42, 11A11, Cell Signalling Technologies; β actin, AC-74, Sigma; GAPDH, 2D9, Origene) in 1% BSA TBST overnight at 4°C before washing and incubation with horseradish peroxidase (HRP) conjugated secondary antibody (1:10,000) (Cell Signalling Technologies). Membranes were then developed using super enhanced chemiluminescence (ECL) (BioRad) and placed in a light proof case and exposed to ECL Hyperfilm (GE Healthcare) for between 1 second and approximately 10 minutes depending on the strength of signal. Chemiluminescence of membranes was also detected using ChemiDoc imager (BioRad) in some instances.

### Intercalation and live cell imaging

Cancer cell TEM or intercalation was determined by live cell imaging. EC were seeded to 96 well plates at a density of 1×10^4^/well in 100μl to achieve confluent monolayers in 24-48 h. Once confluent endothelial monolayers were established, CTG labelled cancer cells were seeded onto endothelial monolayers at a density of 1×10^4^ per well in 50μl of media (total media volume 150μl including endothelial culture medium). Plates were then imaged immediately using Live Cell Imager - Incucyte Zoom. Images were taken every 5 min for 4 h using 20x objective.

### Real Time Cell Analyser (RTCA) analysis using XCELLigence

Real-time monitoring of cellular proliferation, spreading and migration were measured using an electrical impedance-based assay using the xCELLigence RTCA DP instrument (ACEA Biosciences, Inc.). For assessment of spreading and migration, 5×10^3^ MDA-MB-231 cells siRNA reverse transfected (48h post transfection) were seeded to E-plates (ACEA Biosciences, Inc.) and analysed using xCELLigence RTCA DP instrument. Impedance was measured every 15 minutes for 24 h. For real-time monitoring of cellular migration, CIM-plate 16 inserts (ACEA Biosciences Inc.) were used where MDA-MB-231 were plated in serum-free media to the upper chamber and the lower chamber loaded with 10% FCS-containing complete growth media as a chemoattractant. Impedance was measured every 15 minutes for 24 h. Data from each time point were exported and compiled in Excel from a minimum of 3 biological replicates for comparison of siRNA treated cells.

For monitoring of endothelial barrier disruption by cancer cells, E-plates were coated prior to plating endothelial cells, with human fibronectin (Sigma) 10 μg/ml for 2h at RT under a laminar flow hood. Wells were then rinsed twice in PBS and air dried before seeding of endothelial cells. ECs were seeded at 1×10^4^ in 100μl per well of an E-plate and plates were loaded into the xCELLigence RTCA DP instrument. ECs were grown to confluency for 24-48 hours before addition of cancer cells. 1×10^4^ MDA-MB-231 cells in 50μl were loaded to wells containing EC monolayers and impedance was measured every 15 minutes for 24 h. Data were analysed using fold change in barrier integrity upon seeding of cancer cells to EC monolayers compared to HUVEC monolayers alone.

### Extracellular matrix adhesion and immunofluorescence

ViewPlate 96 well black plates (PerkinElmer) were coated with collagen type 1 (10μg/ml in 0.1M acetic acid; Merck Millipore). Plates were incubated at 37 °C for 2 hours before aspirating excess ECM coating and rinsing 2x with PBS and air drying for 30 min at RT. 1×10^4^ cells/well of siRNA treated MDA-MB-231 cells were seeded to coated ViewPlate 96 well plate and allowed to adhere for 30 min. Unbound cells were then washed away in PBS and fixed in 4% PFA for 10 min at RT. Cells were then permeabilised in 0.2% Triton X-100 and blocked for 1 hour in 3% BSA PBS before staining using TexasRed phalloidin (Thermo Scientific (1:200) and Hoechst (1:10,000) in PBLEC staining buffer (0.1% BSA, 1% Tween, 0.05% NaN_3_, 1mM CaCl_2_, 1mM MgCl_2_, 0.1mM MaCl_2_) at RT for 2-4 hours. Following this, cells were rinsed 2x in PBS before subsequent imaging was carried out using an Operetta HTS imager (Perkin Elmer). Image analysis was performed using Columbus software. 3D actin distribution plots were generated using ImageJ ‘3D surface plot’ analysis.

### *In vivo* metastasis model

Female CB17-SCID mice (aged 6-8 weeks) were purchased from Charles River Laboratories. All *in vivo* procedures were performed in accordance with the Animals (Scientific Procedures) Act 1986 Amendment Regulations 2012 following ethical review by the University of Leeds Animal Welfare and Ethical Review Committee (AWERC) Home Office UK PPL No. 70/7544. 72 h following siRNA knockdown, luciferase expressing MDA-MB-231 cells were detached using trypsin and resuspended in RPMI supplemented media. Cells were strained using 70μm cells strainers (Thermo Scientific) to remove aggregates of cells and were pelleted before resuspending at 5×10^6^/ml in PBS. 5×10^5^ cells/mouse were injected into the main tail vein of mice (injections were performed by AT under the approved UK Home Office project license (No. 70/7544)). For imaging, D-luciferin substrate (Sigma) was injected intraperitoneally into mice (2μg/mouse) and mice were anaesthetised with isoflurane and scanned using an IVIS Spectrum imaging unit (PerkinElmer). Bioluminescent imaging of mice was carried out at 6h, 24h, 48h and 72h post injection and then weekly for 4 weeks. For analysis of tumours, formalin-fixed paraffin-embedded lungs were sectioned (6µm), de-waxed and rehydrated prior to staining with H&E or processing for immunofluorescence microscopy following heat-induced antigen retrieval (10mM sodium citrate pH 6.0, 0.1% Tween 20). Sections were stained overnight with rabbit anti-Ki67 AB9260 (Millipore) and mouse anti-CD99 (12E7; BioLegend). Images were captured on an EVOS FL microscope (Life Technologies) and protein expression quantified in ImageJ as outlined previously [23]. Briefly, equivalent captured images were subjected to constant thresholding prior to intensity density measurement. These values were used to calculate corrected fluorescence intensity and percentage area covered.

### CDC42 pulldown

GTP-bound CDC42 was assessed using CDC42 Activation Assay Biochem Kit^TM^ (Cytoskeleton, Inc.). Cells were grown to confluency in T75 flasks, then detached, washed in appropriate media and counted. Cell density was adjusted so equal numbers of cells were used per condition. 3-5×10^6^ cells were reseeded to 0.2% gelatin coated T75 flask at equal densities (per condition of siRNA treatment), and left to adhere for 30 minutes for HUVEC and 4 hours for MDA-MB-231 cells. Following this, cells were placed on ice and media removed. Cells were gently washed in ice cold PBS which was quickly aspirated. Cells were then lysed in 500μl ice cold lysis buffer (provided in kit) and lysates collected using a cell scraper and placed in 1.5ml Eppendorf tubes on ice. 300-800 μg of protein were used (as per the Cdc42 Activation Assay Biochem Kit^TM^ protocol) and added to 20 μl (20 μg) of p21 activated kinase 1 protein (PAK1) – p21 binding domain (PBD) coated beads which were incubated on a rotator at 4°C for 1 h. PAK1-PBD beads and lysate were then pelleted by centrifugation at 3-5,000 x *g* at 4°C for 1 min. Pelleted beads were then mixed with 20 μl 2x Laemmli sample buffer and thoroughly resuspended. Samples were boiled at 100°C for 2 minutes before analysis by SDS-PAGE using 15% gels. Input control lysates, which had not been subject to immunoprecipitation with PAK1-PDB beads were electrophoresed alongside pulldown samples to determine total CDC42 protein expression.

### Coculture tubulogenesis assay

Based upon the method of Bishop et al [24], we seeded HDF at density of 2×10^4^ cells per 0.2% gelatin coated 48 well plate in supplemented ECBM:RPMI media at a ratio of 1:1. Following incubation overnight at 37°C under 5.0% CO_2_, HUVEC cells were seeded at a density of 5-6×10^3^ cells per well of HDFs. Cocultures were cultured for 24 hours before media was replaced with ECMB for controls wells, ECBM + VEGFA (10ng/ml or 25ng/ml)(PeproTech). VEGF supplemented media was changed every 2 days for a period of 9 days before cocultures were washed 2x in PBS and then fixed in 4% PFA for 10 minutes. Wells were washed twice in PBS before staining with a fluorophore conjugated antibody mouse anti-CD31 (Biolegend) (1:400 dilution) in PBLEC staining solution (0.1% BSA, 1% Tween, 0.05% NaN_3_, 1mM CaCl_2_, 1mM MgCl_2_, 0.1mM MaCl_2_) for 1-2 hours at RT with constant agitation, after which cocultures were washed 2x in PBS before imaging using EVOS microscope and Live Cell Imager - IncuCyte.

### Cell proliferation *in vitro*

Cell proliferation was determined using a crystal violet staining assay [25]. Briefly, 1×10^4^ cells were cultured for 24, 48, 72 and 96 hours under normal culture conditions following siRNA transfection. Following culture for indicated time periods cells washed, fixed and stained in 1ml of crystal violet staining solution (0.5% crystal violet powder (Sigma-Aldrich), 20% Methanol) for 20 min at RT with gentle agitation. Stained cells were solubilised in 1ml 100% Methanol/well and incubated with gentle agitation for 20 min at RT with lids on the plate. Optical density of each well was measured at 570 nm (OD_570_) using a plate reader. Average OD_570_ of control empty wells was subtracted from OD_570_ of wells containing cells.

### Scratch wound assay

MDA-MB-231 cells were seeded at a density of 2×10^4^/well of a 96 well plate in 100μl of 10% FCS supplemented RPMI media and grown to confluency over 24-48 hours. Once confluent, cells were serum starved for 2 hours prior to ‘wounding’. Culture medium was removed using an aspirator and replaced with serum free RPMI media (supplemented with 1% BSA). Following serum starvation, scratch wounds were made using the ‘Incucyte wound maker implement’. Once scratched, medium was gently removed by aspirator and monolayers gently washed with PBS before replacing with supplemented RPMI media. The assay was imaged following 18-24 hours to assess closure of the wound. Images were analysed using ImageJ ‘MRI_Wound_Healing_Tool-1’ plugin.

### Patient gene expression data

We downloaded the RNA-seq data generated by Varešlija et al [26], and analysed the expression of VCAM1, SOX9 and CD99 genes in the matched primary and brain tumour metastases from 22 breast cancer patients using paired, non-parametric statistical testing.

### Statistical analysis

Statistical analysis was performed on data generated from n≥3 biological replicates using GraphPad Prism7 software. Error bars represent standard error or standard deviation from the mean as described in the figure legends. The different statistical tests used for individual experiments are also described in the figure legends. Results were deemed significant if p<0.05 and were denoted as: *p<0.05, **p<0.005, ***p<0.0005.

## Results

### Tumour cell CD99 regulates adhesion and transendothelial migration

Leucocytes utilise CD99 to cross endothelial barriers [7–9], suggesting that tumour cell CD99 might regulate this process during metastasis. Cell surface CD99 was readily detected on human breast cancer cell lines (Fig. 1a) and adhesion of fluorescent-labelled MDA-MB-231 cells to an EC monolayer was inhibited by an anti-CD99 antibody (Fig. 1b, c). Endothelial cells also express CD99 (ref. 8) and we used RNA interference (RNAi) to determine whether it was tumour or endothelial CD99 that regulated these adhesion events. Expression of CD99 was reduced in both tumour and EC using siRNA (Fig. 1d and e) and CD99 depleted MDA-MB-231 cells showed significantly reduced adhesion to EC monolayers at early timepoints (Fig. 1f). In contrast, loss of EC CD99 did not significantly decrease tumour cell adhesion and, at a late time point, adhesion was significantly increased, revealing a non-equivalent role for CD99 in the tumour cells and interacting endothelium (Supplementary Figure S1).

**Fig. 1.**
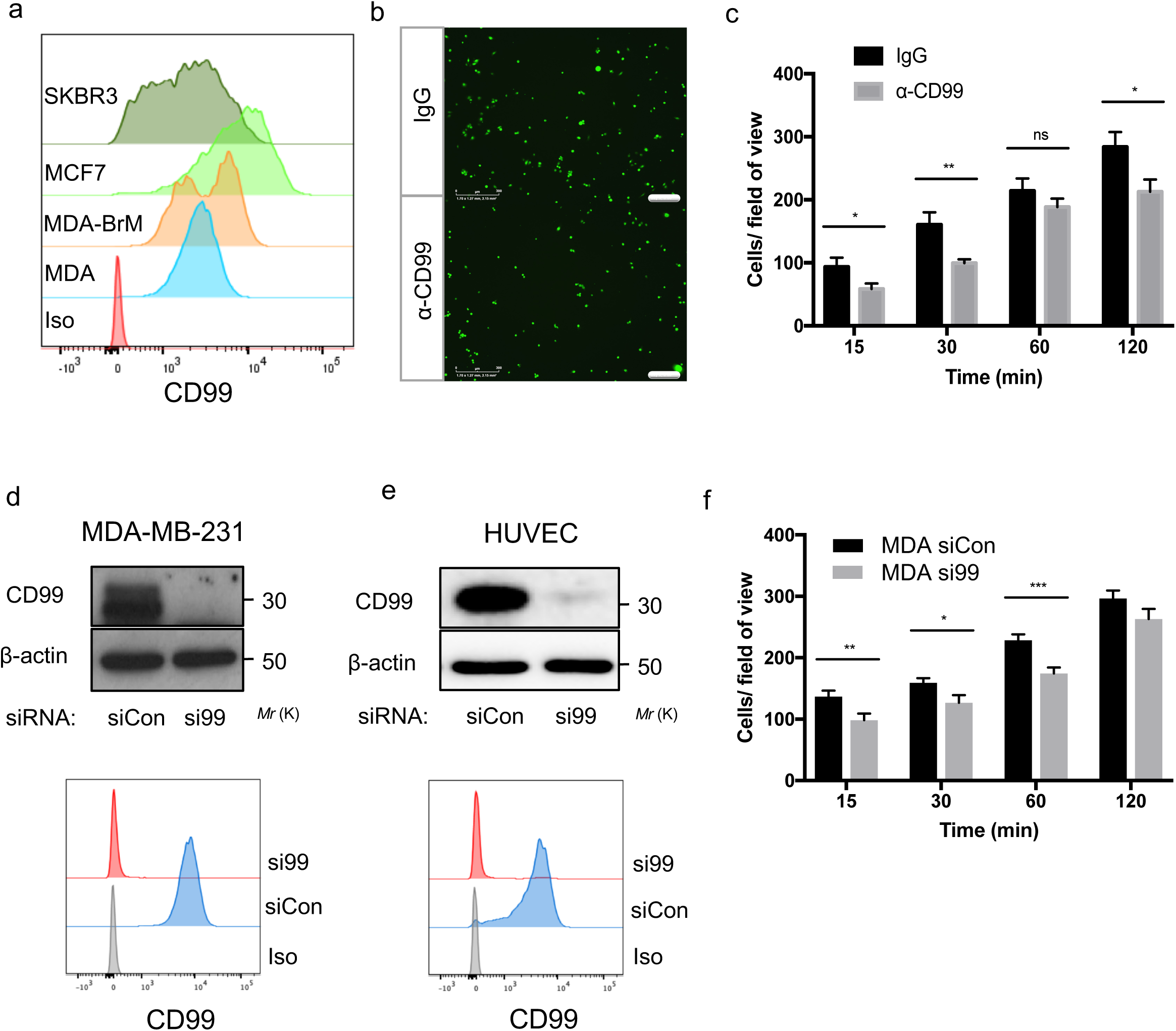
CD99 regulates breast cancer adhesion to endothelial cells. **a** Expression of cell surface CD99 on breast cancer cell lines SKBR3, MCF7, MDA-MB-231 (MDA) and MDA-MB-231 BrM (MDA-Brm, a variant which preferentially metastasises to brain) determined by flow cytometry compared to isotype control (Iso). **b** MDA-231 cells were Cell Tracker Green (CTG)-labelled and left to adhere (for indicated time) to confluent HUVEC monolayers pre-treated with anti-CD99 blocking antibody (1μg/ml) or an IgG control. Unbound cells were washed away with PBS before fixation in 4% PFA and quantification of bound cells. Scale bar: 300 μm. **b** shows representative images with quantification in **c** (using ImageJ) of bound MDA-MB-231 cells. **d** siRNA mediated knockdown of CD99 determined by western blotting and flow cytometry in MDA-MB-231 and **e,** HUVECs using anti-CD99 antibody and anti-β actin as a loading control. As in **a,** flow cytometric analysis of surface CD99 compared to isotype control (iso)**. f** MDA-MB-231 cells were transfected as in **d**, for 72h before CTG-labelled and left to adhere to confluent HUVEC monolayers for indicated time points, as in **b** and **c**. Unbound cells were washed away with PBS before fixation, imaging and quantification of bound MDA-MB-231 cells using ImageJ. Values represent mean±s.e.m., *p<0.05, **p<0.005, ***p<0.0005, Student’s t-test, n=3 independent experiments.

Following adhesion, extravasating cells undergo TEM by crawling between, or sometimes through, endothelial cells [3]. Time-lapse microscopy was used to monitor the interactions of fluorescent-labelled MDA-MB-231 cells with an unlabelled EC monolayer (Fig. 2). Tumour cells interacting with the endothelium altered their morphology from spherical structures towards a spread, more flattened appearance and depletion of tumour cell CD99 increased this spreading phenotype, an effect which was statistically significant at four-hours (Fig. 2a, b). This phenotype was also observed using the metastatic prostate cancer cell line PC3 (Supplementary Figure S2). In the breast cancer model, approximately 30% of MDA-MB-231 cells depleted of CD99 were flattened onto the endothelial monolayer after four hours, compared to approximately 20% of the cells treated with control siRNA (Fig. 2b). Furthermore, increased spreading in CD99 depleted cells was independent of interaction with EC, with increased spreading also occurring on both gelatin and collagen A matrices (Fig. 2c).

**Fig. 2.**
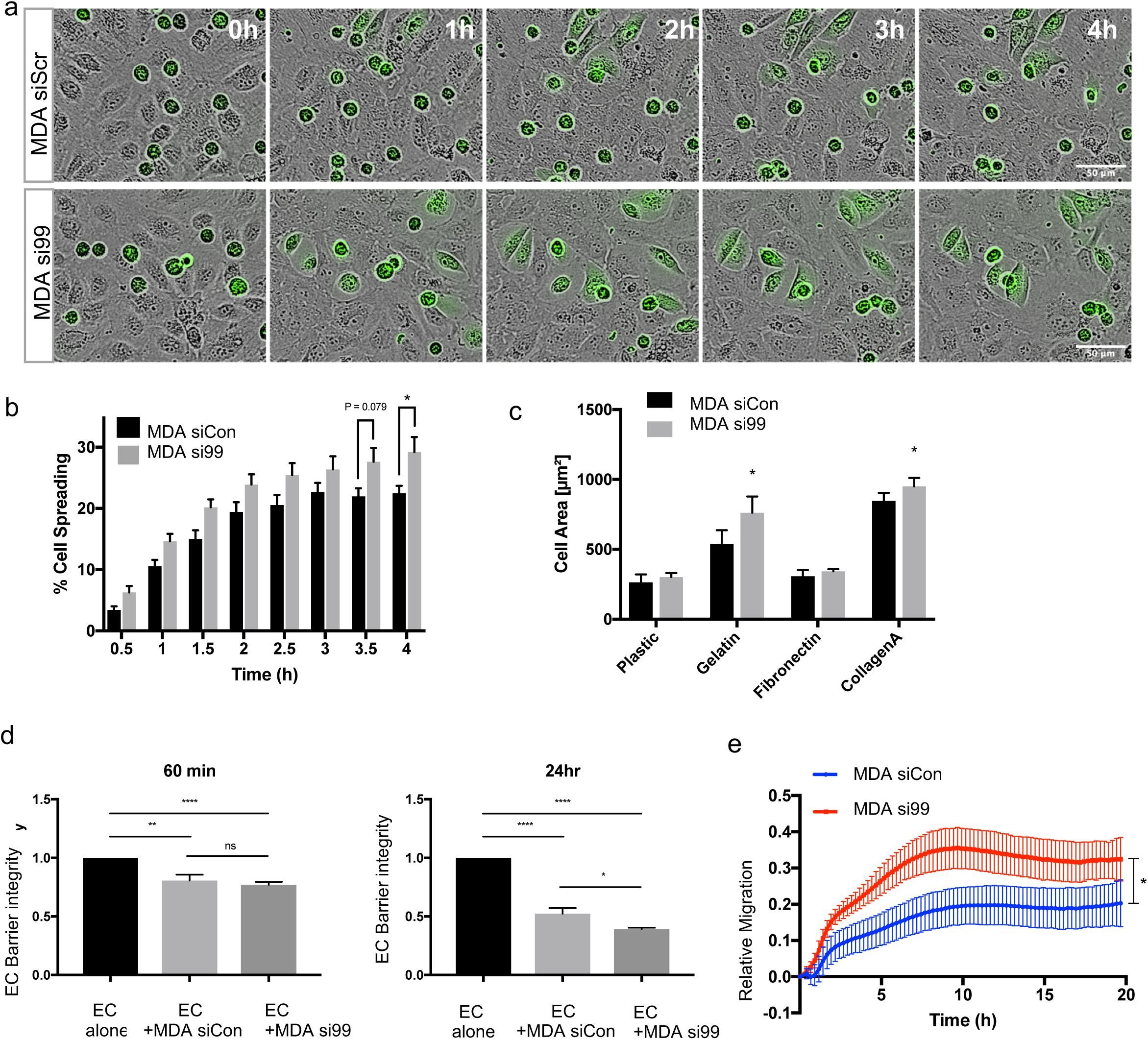
Breast cancer migration and invasion is regulated by CD99. **a** MDA-MB-231 TEM and intercalation determined by live cell imaging. Control or CD99 siRNA treated MDA-MB-231 were CTG-labelled and seeded onto HUVEC monolayers and intercalation/spreading captured using live cell imaging. Images were taken every 5 min for 4 h using 20x objective. Scale bar: 50µm. **b** Quantification of data in panel **a**, indicating percentage of MDA-MB-231 which have undergone spreading/intercalation as a percentage of total cells. Values represent mean±s.e.m., *p<0.05, Student’s t-test, n=3 independent experiments. **c** Cell area of CD99 siRNA transfected MDA-MB-231 cells bound to indicated ECM components for 30 min was determined using Columbus image analysis software. Values represents mean±s.d., *p<0.05, Student’s t-test, n=3 independent experiments. **d** MDA-MB-231 cells treated with control or CD99 siRNA were seeded onto confluent HUVEC monolayers and changes in impedance were recorded at 15 min intervals for 24h at 37°C under 5% CO_2_. Data is presented as fold change in barrier integrity compared to HUVEC monolayers alone at indicated time points. **e** Migration of CD99 siRNA-treated MDA-MB-231 cells determined using the modified Boyden chamber Cell Invasion/Migration (CIM) plates and xCelligence Real Time Cell Analyser (RTCA). Changes in impedance are indicative of the number of cells that have migrated. Impedance was measured every 15 minutes for 24h. Graph represents mean±s.e.m., ***p<0.0005, Analysis of Covariance (ANCOVA), n=4 independent experiments.

Tumour cell invasion of an endothelial barrier requires EC-EC junctions to be broken and the resultant loss in monolayer integrity can be assessed by measuring the reduction in electrical resistance across the monolayer [27]. Addition of tumour cells to an EC monolayer resulted in a significant loss of endothelial barrier integrity at both one hour and twenty-four hours, consistent with TEM activity (Fig. 2d). Tumour-mediated loss of barrier integrity was independent of CD99 expression at the one-hour timepoint. However, CD99 depleted MDA-MB-231 cells displayed a significantly enhanced ability to disrupt the endothelial barrier at the twenty-four hour timepoint (Fig. 2d).

The ability of inflammatory cells or tumour cells to intercalate and cross the endothelium is a migratory function (hence the term TEM). End-point migration assays (using scratch wound repair) showed a trend towards greater migratory activity in the absence of CD99 (Supplementary Figure S3). Furthermore, in real time migration assays (using RTCA modified Boyden chambers), CD99 depletion significantly increased migratory activity of MDA-MB-231 cells (Fig. 2e), consistent with the inhibitory effects of tumour cell CD99 expression on intercalation, cell spreading and endothelial barrier function (Fig. 2a-d).

### CD99 regulates cytoskeletal dynamics and CDC42 activity

Leucocyte extravasation through endothelial monolayers requires substantial cytoskeletal remodelling [28] and similar mechanisms operate in extravasating cancer cells [5]. We plated RNAi-treated MDA-MB-231 cells onto collagen A (where the enhanced spreading phenotype of CD99-depleted cells mirrored that found on the endothelial cells; Figures 2a-c) and analysed their actin organisation using immunofluorescence microscopy; the flattened morphology of the CD99 depleted cells was accompanied by actin rearrangement, with its accumulation at the cell periphery (Figure 3a). The ability of cell surface CD99 to regulate actin dynamics suggested a potential role for the Rho GTPase, CDC42. Active CDC42 (with bound GTP) interacts with and activates the serine/threonine kinase PAK1 and hence bead-bound PAK1 can be used to selectively pull down active CDC42, which can be identified using anti-CDC42 antibodies [29]. Using this approach, we found that CD99 depletion in MDA-MB-231 cells resulted in an increase in active CDC42 (Figure 3b); interestingly levels of total CDC42 also seemed to be increased under these conditions (Figure 3b). These results are consistent with a pathway in which cell surface CD99 regulates cytoskeletal dynamics via alterations in CDC42 activity; specifically, CD99 expression negatively regulates CDC42 activity.

**Fig. 3.**
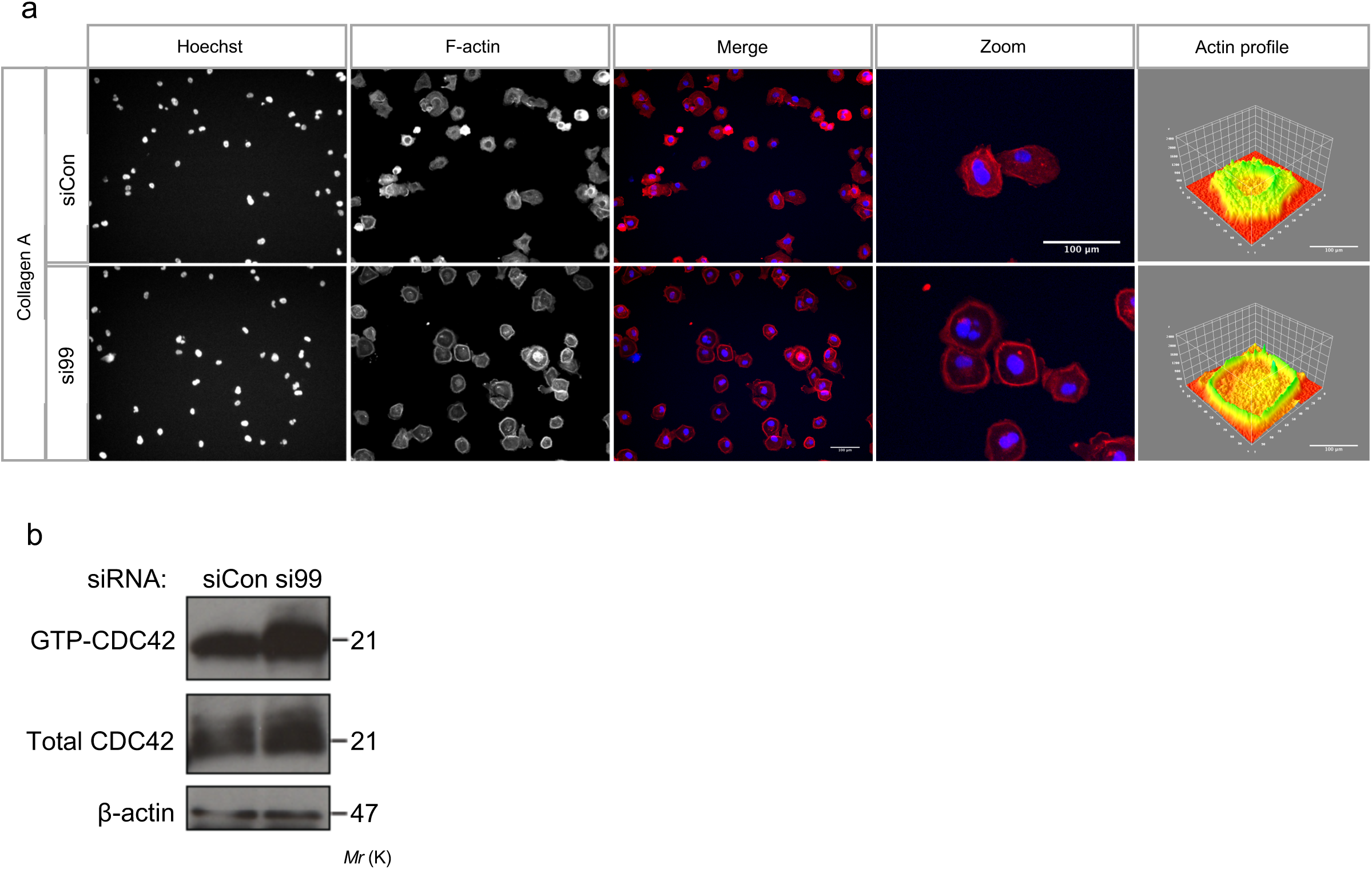
Breast cancer actin dynamics are modulated by negative regulation of CDC42 by CD99. **a** Scrambled or CD99 siRNA treated MDA-MB-231 cells (48 hr post transfection) were re-plated to collagen type 1 coated plates and allowed to adhere for 30 min. Cells were then stained with TexasRed phalloidin before imaging using an Operetta HTS imager. 3D actin distribution plots were generated using ImageJ ‘3D surface plot’ analysis. Scale bar: 100 µm. Images are representative of n=3 independent experiments. **b,** MDA-MB-231 were transfected with scramble control and CD99 siRNA and lysates were subject to immunoprecipitation with PAK1-PDB beads and subsequent electrophoresis to determine GTP-bound CDC42. Input control lysates, which had not been subject to immunoprecipitation with PAK1-PDB beads were electrophoresed alongside pulldown samples to determine total CDC42 protein expression.

### Tumour cell CD99 negatively regulates metastasis *in vivo*

These results highlight a complex role for CD99 in tumour cell TEM; CD99 is coupled to cytoskeletal organisation and positively regulates tumour-EC adhesion, but negatively regulates post-adhesion events, including cell migration. Such complexity reflects the multistage nature of tumour cell TEM and is consistent with previous findings suggesting either pro- or anti-tumour activity for CD99 [11]. To determine the overall contribution of CD99 expression to metastasis and tumour progression we used an *in vivo* model. When injected into the tail vein of immunodeficient mice, the MDA-MB-231 cell line preferentially metastasises to the lungs. We monitored progression of control and CD99 depleted, luciferase expressing MDA-MB-231 cells over a four-week period and found that CD99 depleted MDA-MB-231 cells exhibited greater metastatic activity than the control siRNA treated cells (Fig. 4a, b). Upon histological analysis, the lung metastases in the CD99-depleted MDA-MB-231 cells were more densely packed than metastases seeded by control treated cells (Figure 4c), consistent with the increased luciferase signal from *in vivo* imaging. Within tumours, a significantly greater proportion of cells from the si99 treated group expressed ki67, consistent with the increased tumour burden. However, overall ki67 expression levels were not significantly different between the control of si99 treated groups, indicating similar levels of proliferation (Figure 4e). Furthermore, tumour cell CD99 did not significantly affect proliferation *in vitro* (Supplementary Figure S4). This xenograft model identifies tumour cell CD99 as an inhibitor of tumour progression, an effect that is independent of proliferation. To analyse CD99 expression in patients, we made use of published RNA-seq data obtained from twenty-three primary breast cancers and their matched brain metastases [26]. A number of genes have previously been shown to be differentially expressed between the primary tumour and the brain metastases. For example, VCAM1 is downregulated in brain metastases, whereas SOX2 is upregulated [30]. We confirmed this pattern of expression in the data of Varešlija et al [26] and further analysis showed that CD99 mRNA was significantly downregulated in the metastases (Figure 4f).

**Fig. 4.**
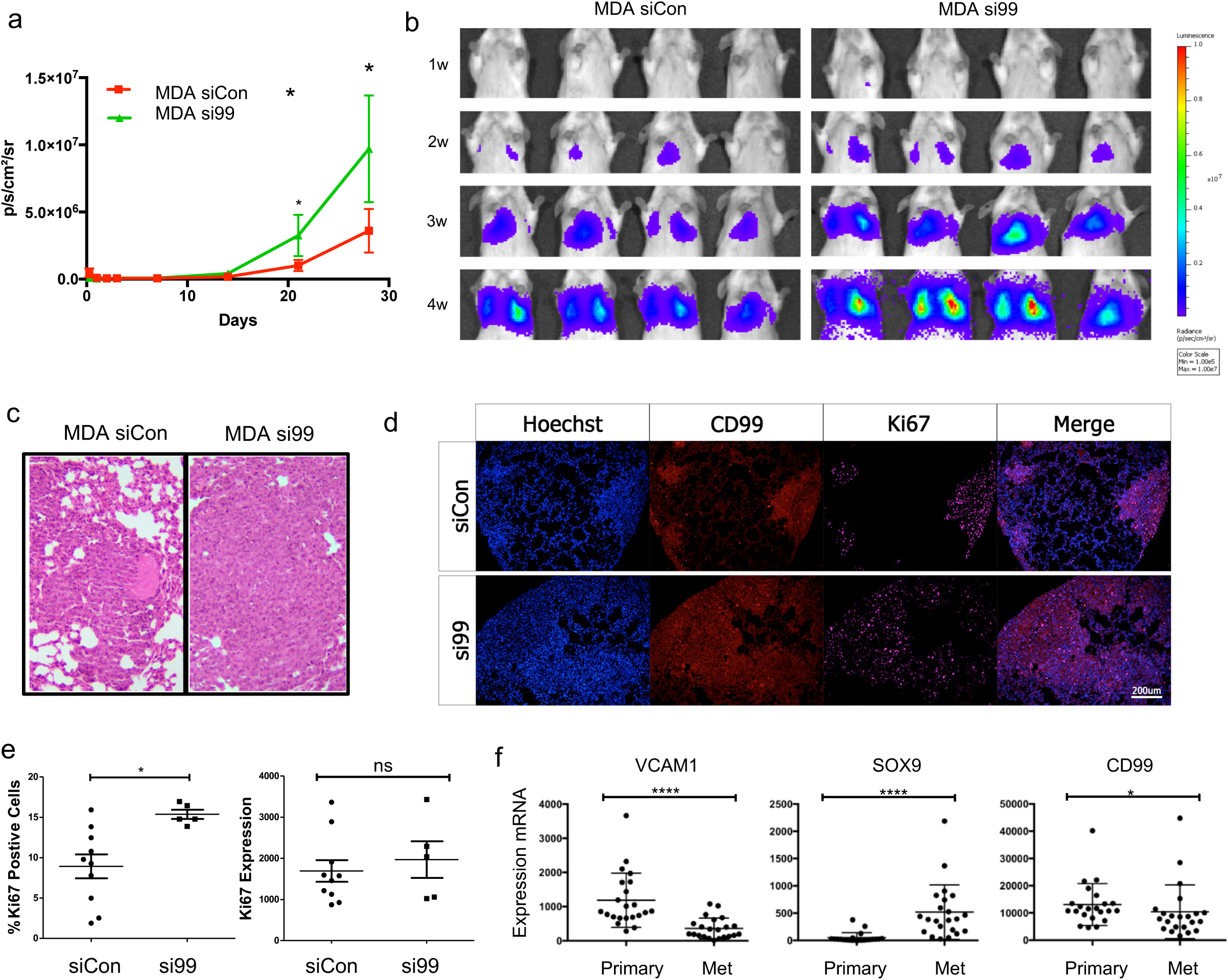
CD99 suppresses the metastatic phenotype of breast cancer *in vivo.* Luciferase expressing MDA-MB-231 cells were transfected with scramble control or CD99 siRNA. 72 h following siRNA knockdown, 5×10^5^ cells/mouse were injected into the main tail vein of female CB17-SCID mice (aged 6-8 weeks). **a** Quantification of bioluminescent imaging from each time point. Values represent mean±s.d, *p<0.05, Student’s t-test, n=4 per group. **b,** For imaging, D-luciferin substrate was injected and anaesthetised mice scanned using IVIS Spectrum Imaging. **c,** Representative images of lung tissue from mice injected with control or CD99 siRNA treated MDA-MB-231 cells. Lungs were isolated 4 weeks post injection and analysed by H&E staining. d, Lung sections stained with Hoechst (to detect all cells), anti-CD99 antibody (to detect the human tumour cells) and anti-ki67 antibody (to detect proliferating cells); Scale bar: 200µm. e, Quantification (using ImageJ) of ki67 staining as a percentage of total Hoechst nuclei (left panel) and relative fluorescence intensity (right panel). Data was analysed using an unpaired Student’s t test; *p<0.05. **f,** Gene expression in primary and matched brain tumour metastases from 22 breast cancer patients (25); data was analysed using a Wilcoxon matched pairs signed rank test; *p<0.05, ****p<0.00005.

### EC CD99 negatively regulates CDC42 and downstream angiogenic activity

Transendothelial migration requires cytoskeletal remodelling of both the transmigrating cell (the leucocyte or tumour cell) and the endothelial cells [3–5]. The expression of CD99 on the endothelium (Ref. 8 and Figure 1e) and the proven role of endothelial cell CD99 in leukocyte TEM [7–9] suggests that endothelial cell CD99 might also regulate tumour cell TEM. Antibody blocking of CD99 arrests migrating leucocytes partway through the TEM process [7]. We observed a similar phenotype, whereby CD99 depleted EC monolayers reduced the ability of MDA-MB-231 cells to spread and penetrate the endothelial barrier (Fig. 5a). Quantification showed that CD99 depletion significantly delayed the intercalation of the breast cancer cells, however this effect was abrogated at later time points (Fig. 5b); similar results were obtained using siRNA-treated ECs and prostate cancer PC3 (Supplementary Figure S5). As previously shown in Fig. 2d, addition of MDA-MB-231 cells to an EC monolayer disrupted barrier function (Fig. 5c). However, whilst barrier disruption was evident at 30 minutes using control EC, disruption of CD99 depleted EC was slower (detectable at 90 minutes; Fig. 5c), consistent with the reduced spreading of tumour cells onto CD99 depleted EC monolayers seen in Fig. 5a and b. Thus, EC CD99 promotes intercalation/TEM of tumour cells, whereas tumour cell CD99 negatively regulates TEM (Fig. 2d).

**Fig. 5.**
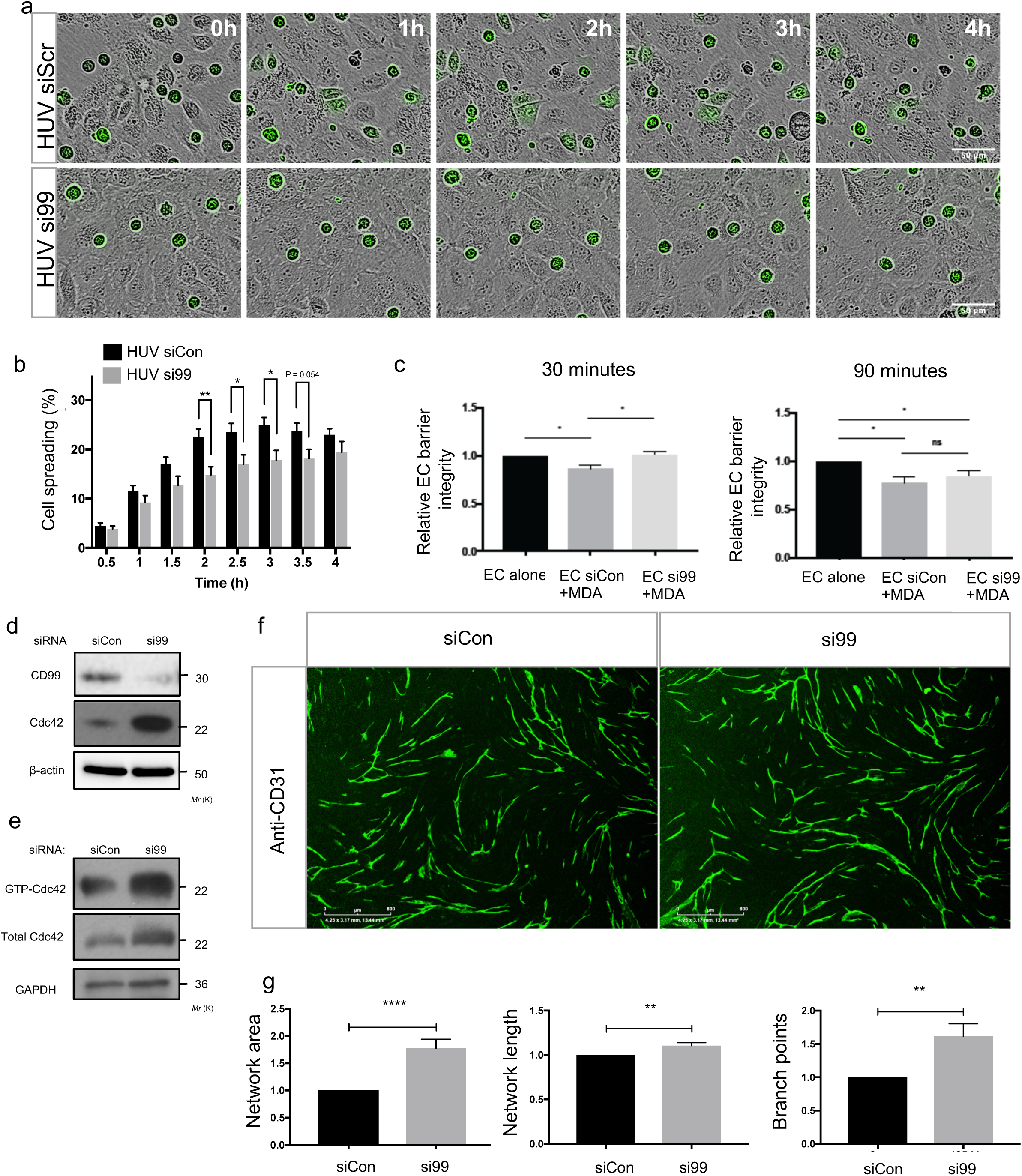
Endothelial CD99 regulates cancer transendothelial migration and angiogenesis by modulating CDC42 activity and expression. **a** MDA-MB-231 transendothelial migration and intercalation was determined by live cell imaging. CTG labelled MDA-MB-231 were seeded to Scrambled or CD99 siRNA treated HUVEC monolayers and imaged using live cell imaging to capture MDA-MB-231 spreading. Images were taken every 5 min for 4 h at 20x objective. Scale bar: 50 µm. **b** Quantification of data in panel **a**, showing spreading of MDA-MB-231 cells as a percentage of total cells, indicating percentage of MDA-MB-231 which have undergone transendothelial migration. Values represent mean±s.e.m., *p<0.05, **p<0.005, Student’s t-test, n=3 independent experiments. **c** HUVEC were transfected with CD99 or scrambled siRNA were seeded to XCelligence E-plates to monitor barrier integrity of monolayers. MDA-MB-231 cells were seeded on top of HUVEC monolayers and changes in impedance were recorded. Data is presented as fold change in barrier integrity compared to HUVEC monolayers alone. Values represent mean±s.e.m., *p<0.05, Student’s t-test, n=3 independent experiments. **d** Scrambled or CD99 siRNA treated HUVEC were analysed by western blot for CD99, CDC42 and β actin expression levels. **e** HUVEC were transfected with scramble control and CD99 siRNA and lysates were subject to immunoprecipitation with PAK1-PDB beads and subsequent electrophoresis to determine GTP-bound CDC42. Input control lysates, which had not been subject to immunoprecipitation with PAK1-PDB beads, were electrophoresed alongside pulldown samples to determine total CDC42 protein expression. **f** Representative images of siRNA treated HUVEC tubules. Scrambled or CD99 siRNA treated HUVEC were cocultured with human dermal fibroblasts (HDF) for 7 days and stimulated with VEGFA (10ng/ml) before fixation and staining for CD31. Scale bar: 800μm. **g** Quantification of network area, length and branch points were analysed using ImageJ. Values represents mean±s.e.m., *p<0.05, **p<0.005, ***p<0.0005, ****p<0.00005, Student’s t-test, n=3 independent experiments.

Endothelial barrier integrity is in part regulated by CDC42 via maintenance of adherens junctions at EC-EC contacts [16, 17, 26]. We investigated whether the functional coupling of CD99 to CDC42 activity evident in MDA-MB-231 cells (Figure 3b) was also evident in EC; RNAi-mediated depletion of CD99 in EC resulted in greater expression of total CDC42 (Fig. 5d) matched by an increase in the active, GTP bound CDC42 (Fig. 5e). Thus, in EC, CD99 negatively regulates CDC42 expression and activity, thereby regulating EC cytoskeletal dynamics and barrier function. These results suggest that EC CD99 facilitates TEM of tumour cells by negatively regulating CDC42 activity, consistent with the functional coupling of cell surface CD99 to CDC42 observed in tumour cells.

The CDC42 molecule has previously been shown to play a key role in EC angiogenic activity [32][33][34], suggesting that CD99 might also regulate angiogenesis via its regulation of CDC42 activity. Angiogenesis requires migration and proliferation of EC [35], events that are accompanied by CDC42-dependent cytoskeletal remodelling [18, 28, 29]. This suggested that EC CD99 might regulate angiogenic activity. Angiogenic potential can be assayed by co-culturing EC with fibroblasts in the presence of VEGFA; we compared normal and CD99 depleted HUVEC for their ability to form vascular networks and tubules. CD99 depletion enhanced the network length, the number of branch points and total network area over a seven-day period (Fig. 4g, h), consistent with the increased in active CDC42 in the CD99 depleted HUVECs (Fig. 5e). Thus, EC CD99 negatively regulates CDC42 activity and downstream angiogenic responses. By promoting cytoskeletal rearrangement via CDC42 activation, the loss of EC CD99 favours the migratory activity of EC required for angiogenesis.

## Discussion

Metastasis is a complex, multistep process involving migration, invasion and accompanying changes in the actin cytoskeleton [1, 3]. Our results identify a functional link between CD99, CDC42 and cytoskeletal dynamics in both cancer cells and endothelial cells and demonstrate how CD99 regulates TEM pathways in both the endothelium and the transmigrating cell type. Previous studies have linked CD99 engagement to actin reorganisation [36]. Furthermore, the CD99 cytoplasmic tail associates with a complex of sAC, PKA and ezrin [9], the latter of which is an ERM protein linking the plasmamembrane to the cytoskeleton [37]. Interestingly, CDC42 is activated by both cAMP and PKA [38], implicating CD99-associated sAC and PKA in CDC42 regulation. Our data showing that depletion of tumour cell CD99 increases CDC42 activity and enhances the spreading and intercalation of tumour cells is consistent with the work of Reymond *et al* showing that CDC42 depletion reduced both tumour cell spreading on endothelial monolayers and the metastasis of MDA-MB-231 cells *in vivo*; this study also showed that CDC42 positively regulated β1 integrin gene expression [20]. Increased angiogenic activity upon CD99 depletion in EC may be linked to CDC42-mediated changes in integrin expression [34–36]. Furthermore, β1 integrin null mice exhibit perturbed EC polarity coupled with redistribution of junctional molecules, including CD99 [42]. Indeed, CDC42 activity modulates numerous cellular pathways in response to diverse stimuli [15, 43] and the ability of CD99 to modulate CDC42 activity suggests that CD99 will similarly regulate multiple events. CDC42 has well documented roles in angiogenesis and endothelial cell function and the increased angiogenic activity of EC we observed upon CD99 depletion is likely to be a combination of CDC42-mediated actin remodelling coupled with additional CDC42 regulated pathways known to impact on angiogenesis [16–18,20,32–34].

Expression studies reveal that CD99 has both positive and negative effects on human tumour progression, depending on the tumour type [11]. For example, high expression of CD99 in Ewing’s sarcoma favours migratory activity *in vitro* and RNAi-mediated depletion of CD99 reduces metastasis in a xenograft model [12]. Interestingly, loss of CD99 in Ewing’s sarcoma resulted in neurite outgrowth, a process shown in other cells to require cytoskeletal rearrangements and Rho GTPases, including CDC42 [44–46]. Similarly, CD99 is highly expressed in various brain tumours and, in glioma, reduced expression was associated with decreased migration and altered Rho and Rac activity [11, 47], further supporting functional linkage of CD99 to Rho GTPase activity. Such findings imply that CD99 has oncogene-like activity. However, in other cancers CD99 exhibits tumour suppressor-like behaviour; CD99 expression is lost during osteosarcoma development and ectopic expression suppresses migration *in vitro* and tumour progression and metastasis in a xenograft model [48]. Similarly, decreased expression of CD99 in gastric cancer is associated with poor prognosis and the RNAi-mediated CD99 depletion in a gastric cancer cell line resulted in increased migratory activity [49]. It has been reported that CD99 expression is not prognostic in primary breast cancer, however, CD99 expressing tumours were present in low numbers in this study [50].

Documented pro- and anti-tumour activity of CD99 is mirrored by both positive and negative roles for CDC42 in tumour progression [51]. Increased expression of CDC42 is observed in several cancers, including breast and colorectal cancer [52, 53], and in melanoma, lung and testicular cancer, increased expression is associated with poor outcome [54–56]. Furthermore, CDC42 is required for Ras-mediated oncogenesis in mice [57, 58]. However, tumour suppressor-like activity of CDC42 has also been reported, in human cancer as well as mouse models [59–61]. Our data shows that CD99 depleted breast cancer cells exhibit enhanced tumour progression in a metastatic xenograft model and enhanced TEM activity *in vitro*, consistent with tumour-suppressor-like activity of CD99 as suggested by osteosarcoma and gastric cancer [48, 49]. This activity also appears supported by the reduced expression of CD99 mRNA in breast cancer brain metastases compared to their matched primary tumours. However, our xenograft model only represents the later stages of haematogenous metastasis, since the tumour cells were injected directly into the circulation; how CD99 depleted tumour cells behave during the progression of the primary tumour and its intravasation and spread are unclear. Increased progression of CD99 depleted tumour cells is consistent with increased migration in the absence of CD99 and the enhanced ability of CD99 depleted cells to disrupt endothelial cell barriers. However, we cannot dissociate TEM activity in this model from events occurring after seeding into the metastatic site. Proliferation was unaffected by CD99 loss, but tumour-stromal interactions, such as tumour-fibroblast interactions, angiogenesis and innate immune recognition (which remains intact in the SCID mouse model) may also be regulated by tumour cell CD99 and contribute to progression. Interestingly, tumour cell CDC42 has previously been shown to confer resistance to cytotoxic lymphocytes and, in breast cancer cells, CDC42-dependent actin remodelling at the immune synapse reduces killing by natural killer (NK) cells [62, 63]. Hence, in our xenograft model, it is possible that increased CDC42 activity in CD99 depleted tumour cells provides increased protection against NK cell attack whilst the tumour cells travel in the bloodstream.

In conclusion, we show that the cell surface receptor CD99 modulates tumour cell TEM and metastatic progression and we identify CD99 as a negative regulator of CDC42 activity in both tumour cells and EC. As a signal transduction hub, CDC42 activity impacts upon many of the hallmarks of cancer, including the regulation of cytoskeletal dynamics that accompany the migratory and TEM events that underpin the metastatic process. By establishing a functional link between CD99 and CDC42 in both tumour TEM and angiogenesis, we demonstrate that CD99 plays a key role in tumour progression and implicate cell surface CD99 in controlling the diverse activities regulated by the Rho GTPase CDC42.

## Author’s contributions

The study was designed by AJM, AFO, PFJ and GPC. Most experiments were performed by AJM and AFO. AT assisted with the design and the undertaking of the *in vivo* work and GPC performed expression data mining. Data was analysed and interpreted by all authors. GPC drafted the manuscript with PFJ and AJM. All authors contributed to the writing and approval of the final manuscript. GPC and PFJ made equal contributions to this work.

## Acknowledgments

We are grateful to those colleagues who provided materials for this work and for constructive discussion.

## Abbreviations

ARP2/3: Actin-related protein 2/3 complex
SCID: Severe Combined Immunodeficiency
CD99: Cluster Determinant 99
CDC42: Cell Division Cycle 42
EC: Endothelial Cells
ECBM: Endothelial Cell Basal Medium
FACS: Fluorescence Activated Cell Sorting
GAPS: GTPase activating proteins
GEFs: Guanine nucleotide exchange factors
GTP: Guanosine Triphosphate
HDF: Human Dermal Fibroblasts
HUVEC: Human Umbilical Vein Endothelial Cells
LBRC: Lateral Border Recycling Complex
N-WASP: Neural Wiskott-Aldrich Syndrome Protein
PAK1: p21 Activated Kinase
PDB: p21 Binding Domain
PKA: Protein Kinase A
RNA-seq: RNA sequencing
RNAi: RNA Interference
RTCA: Real Time Cell Analyser
sAC: Soluble Adenyl Cyclase
siRNA: Small Interfering RNA
SOX9: Sry-related HMG box 9
TEM: Transendothelial Migration
VCAM1: Vascular Cell Adhesion Molecule 1
VEGFA: Vascular Endothelial Growth Factor A.

## Supplementary figure legends

**Fig. S1.** Adhesion of MDA-MB-231 cells to CD99 depleted HUVECs. HUVEC cells were transfected with control (siScr) or CD99 siRNA (si99) for 72h and grown to confluent monolayers before seeding of CTG-labelled MDA-MB-231 and left to adhere for the indicated time points. Unbound cells were washed away with PBS before fixation in 4% PFA and quantification of bound cells. Error bars indicate standard Error. Statistical analysis was performed using Student’s t-test. *p<0.05.

**Fig. S2.** CD99 suppresses transendothelial migration of metastatic cancer cells. **a** Prostate cancer PC3 cells were transfected with siRNA targeting CD99 (si99) or a control siRNA (siScr) for 72h before analysis of cell surface CD99 by flow cytometry. Cells were stained with anti-CD99 antibody or an isotype control (iso). Data are representative of three independent replicates. **b** PC3 transfected with CD99 siRNA (si99) or control siRNA (siScr) were labelled with Cell Tracker Green before seeding on top of confluent HUVEC monolayers and imaging for 4 hours using an Incucyte live cell imager. Scale bar: 50 µm. **c** Quantification of data in panel b, showing intercalation of PC3 cells (treated with si99 or siScr as indicated) as a percentage of total PC3 cells. Data is representative of n=3 independent replicates. Error bars indicate standard error. Statistical analysis was performed using student t-test. *p<0.05.

**Fig. S3.** CD99 regulates migration and spreading of breast cancer *in vitro*. **a** Scratch wound migration assay. CD99 depleted (si99) or control siRNA (Scr) treated MDA-231 cells were seeded at equal density to 96 well plates until confluent. Cells were serum starved for 2-3 hours before ‘wounding’ using a Wound Maker tool (Essen Bioscience). Migration of siRNA-treated MDA-MB-231 cells was subsequently monitored using Incucyte live cell imaging (over 18 h) and quantified using Image J. Scale bar: 300 µm. This is a representative image from one of five experiments. **b** Quantification of a, from five independent replicates. Error bars indicate standard deviation. Analysis using the Student’s t test demonstrated that differences in migration between treatments were not statistically significant.

**Fig. S4.** *In vitro* proliferation of MDA-MB-231 after CD99 depletion. MDA-MB-231 cells treated with control (Scr) or CD99 siRNA (si99) were seeded to 24 well tissue culture dishes at the same density. Plates were left for 1,2,3 and 4 days before fixing in 4% PFA and subsequent staining with Crystal Violet. Stained cells were solubilised in 100% methanol and spectrophotometry performed using a plate reader. Results were normalised to stained empty wells and background subtracted. Data are normalised to Scr treated cells at day 1 of the assay and show three replicates. Error bars indicate standard deviation. Statistical analysis performed using Student’s t-test demonstrated that proliferation was not statistically significant between the different treatments.

**Fig. S5.** Endothelial CD99 regulates cancer cell transendothelial migration. Cell Tracker Green-labelled PC3 cells were seeded on top of confluent HUVEC monolayers that were transfected with si99 or Scr siRNA and imaged for 4h using live cell imaging via the Incucyte microscope to capture intercalation (as in Figures 2a, 4a and S2b). Quantification of PC3 cell spreading was performed and is displayed as a percentage of total cells. Error bars indicate standard error. Statistical analysis was performed using student t-test. *p<0.05.

