## Supplementary Figures S1-5 for "CD99 regulates cancer cell transendothelial migration and endothelial cell function via CDC42 and actin remodelling"

### Supplementary Figure S1

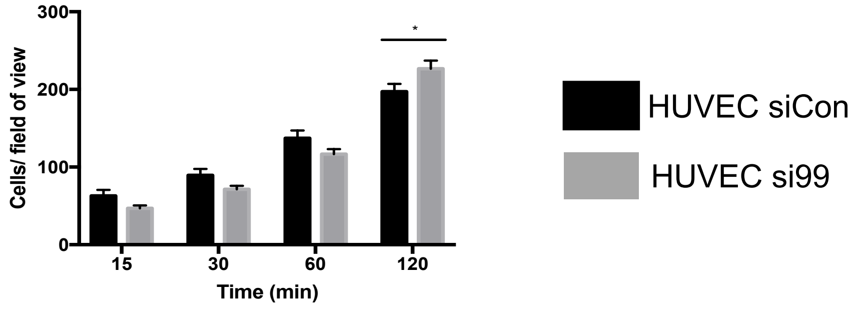


### Supplementary Figure S2

a

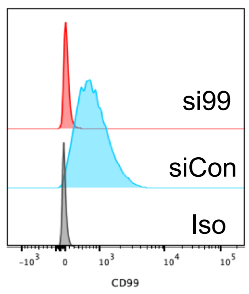


b

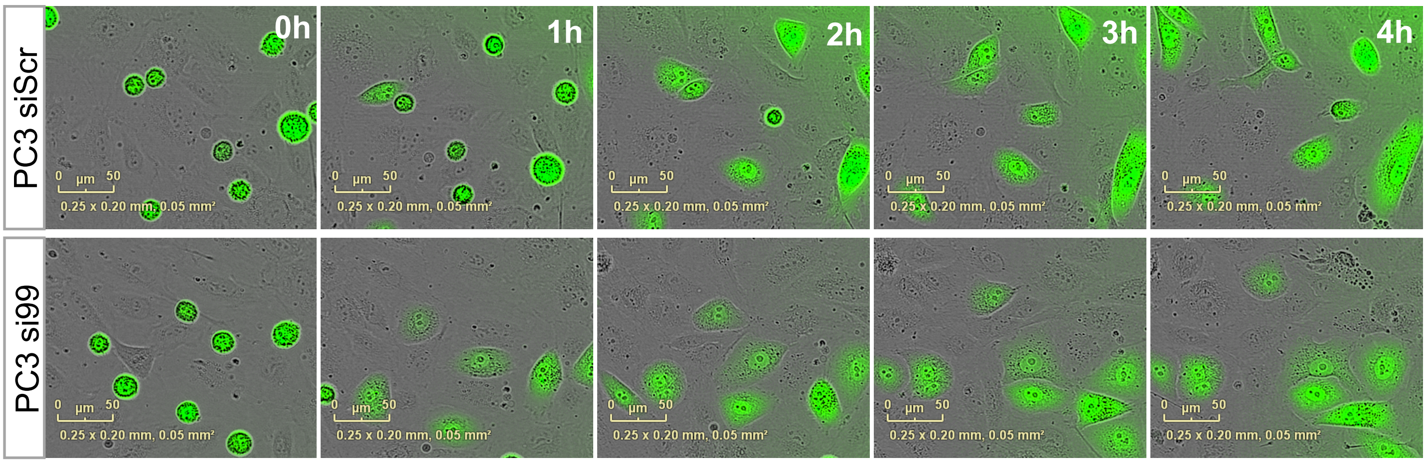


c

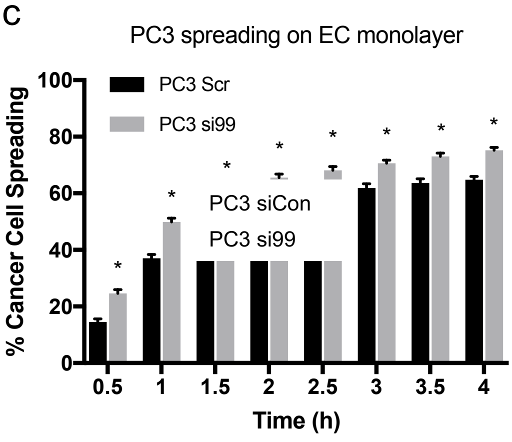

### Supplementary Figure S3

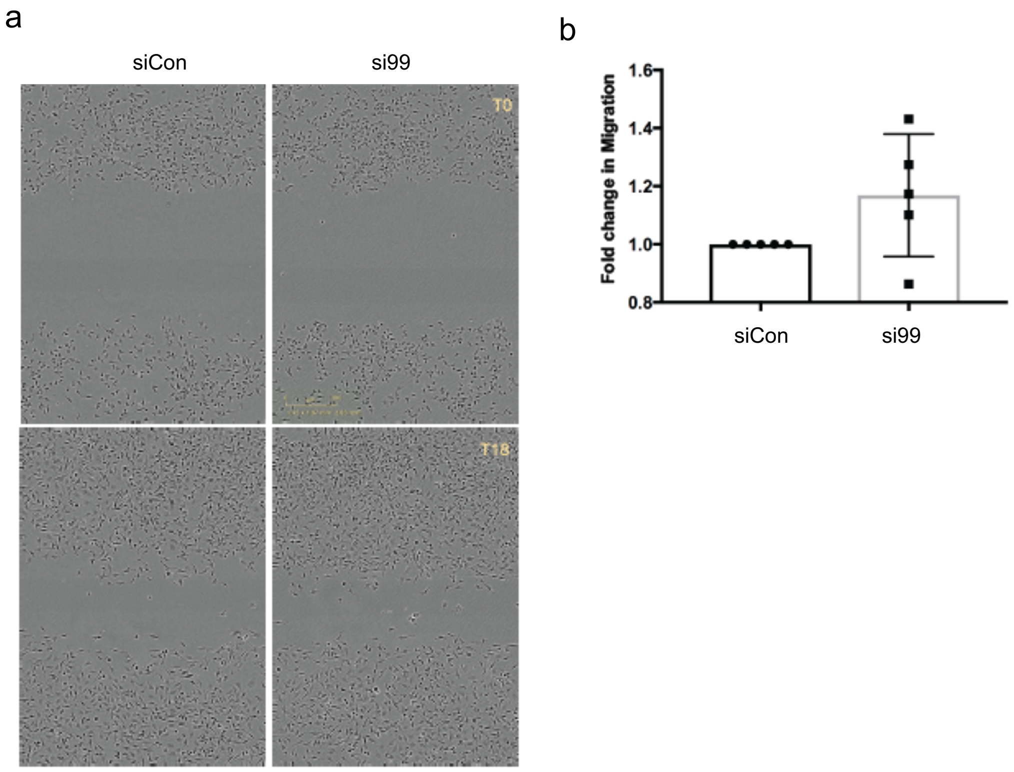


### Supplementary Figure S4

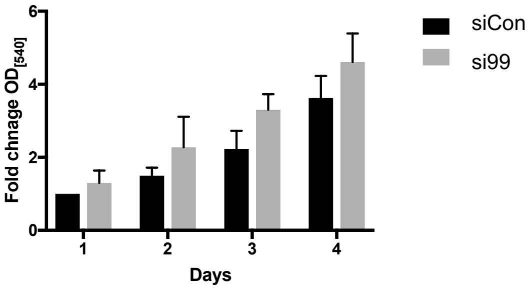


### Supplementary Figure S5

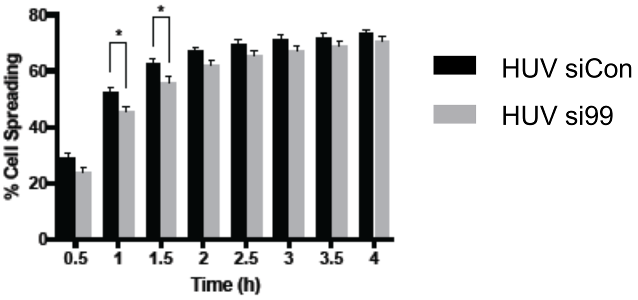
